# Multiple ligand recognition sites in free fatty acid receptor 2 (FFAR2) direct distinct neutrophil activation patterns

**DOI:** 10.1101/2020.10.27.356923

**Authors:** Simon Lind, André Holdfeldt, Jonas Mårtensson, Kenneth L. Granberg, Huamei Forsman, Claes Dahlgren

## Abstract

Non-activating positive allosteric modulators specific for free fatty acid receptor 2 (FFAR2) increased the activity induced by orthosteric agonists to trigger a rise in intracellular Ca^2+^ ([Ca^2+^]_i_) and activate the O_2_^−^ producing neutrophil NADPH-oxidase. In addition, two allosteric modulators (Cmp58 and AZ1729) recognized by different receptor domains on FFAR2, cooperatively triggered activation without any rise in [Ca^2+^]_i_. To gain insights into FFAR2 modulation and signaling, we set out to identify structurally diverse allosteric FFAR2 modulators. Initially, we identified two molecules that directly activate neutrophils and these were classified as an allosteric FFAR2 agonists and an orthosteric agonist, respectively. Based on the sensitizing effect on the neutrophil response to propionate, ten non-direct-activating molecules were classified as allosteric FFAR2 modulators. One of these synergistically activated neutrophils when combined with AZ1729, but not when combined with Cmp58. The remaining nine compounds synergistically induced the same type of biased neutrophil signaling but only when combined with Cmp58. The activation signals down-stream of FFAR2 when stimulated by two allosteric modulators with different binding sites were in most cases biased in that two complementary modulators together triggered an activation of the NADPH-oxidase, but no increase in [Ca^2+^]_i_. The neutrophil activation pattern achieved when two functionally “AZ1729- or “Cmp58-like” allosteric FFAR2 modulators were combined, supporting a model for activation in which FFAR2 has two different sites that selectively bind allosteric modulators. The novel neutrophil activation patterns and receptor down-stream signaling mediated by two cross-sensitizing allosteric modulators represent a new regulatory mechanism that controls FFAR2 receptor function.

## Introduction

The G protein-coupled receptors (GPCRs) comprise a family of cell surface receptors expressed in many different cell types. These receptors have a common structure with a peptide chain that transverse the plasma membrane seven times, leaving the N-terminus on the extracellular side of the membrane and the C-terminal tail facing the cytoplasm. A binding site that recognizes orthosteric agonists is reachable for ligands on the surface of the receptor-expressing cells, and binding of such an agonist to the orthosteric site induces conformational changes of the receptor (1,2). This binding might result in a shift of the binding affinity for the agonists, but more importantly, the agonist induced structural changes affects the cytosolic signal transducing parts of the receptor. This results in an activation of signaling pathways down-stream of the agonist occupied receptor, in order to regulate the functional response of the receptor expressing cells. Signaling adapted to an on/off mechanism determined by whether an agonist is bound to its receptor or not, was for many years the accepted model for how GPCR signaling was thought to be regulated, but it is now known that receptor signaling is more variable. Consequently, the classification of the ligands that interact and regulate receptor functions has been adjusted (3,4); binding of classical GPCR agonists recognized by the orthosteric receptor site may amplify multiple signaling pathways and down-stream responses, whereas non-classical agonists give rise to biased signaling and functionally selective responses (3). The diversity of biased/functional selective responses are illustrative for the variability of receptor activities induced by different agonists that stabilize receptor conformations that promotes a balanced biological response or one signaling pathway over another (3). Receptor selective/specific ligands may also bind to sites that are separated structurally from the orthosteric binding site. Commonly, such ligands are non-activating on their own, yet they modulate receptor signaling and function of orthosteric agonists (5). A receptor with a bound modulating ligand is transferred to an allosteric modulated state that has a new signaling potential (increased or decreased) when activated by a second agonist. Such an allosteric GPCR modulation is, according to the dogma for allosteric GPCR-modulation, solely affecting the signaling properties of agonists that specifically interact with the receptor that is allosterically modulated (6,7). This GPCR-modulation dogma has, however, been challenged and shown to be more complex than initially anticipated, when it was shown that allosteric modulators specific for Free Fatty Acid Receptor 2 (FFAR2) affect signaling mediated not only by orthosteric FFAR2 agonists but also by specific agonists for the ATP-receptor P2Y2R and for formyl peptide receptors (FPRs; (8–10).

The FFARs recognize endogenous orthosteric agonist such as acetate and propionate, that intertwine metabolism and immune function in both health and disease (11) suggesting that new therapeutics that regulate receptor functions may be developed. Accordingly, a novel allosteric FFAR2 ligand, AZ1729, was recently described and shown to be both a positive allosteric FFAR2 modulator that increased the activity induced by conventional orthosteric agonists and a direct activating allosteric agonist (12). It is interesting to note, that the allosterically modulated FFAR2s were able to signal through different G-proteins containing either Gai or Gaq subunits (“induced-bias” (12,13)). Based on earlier findings, it is reasonable to assume that FFAR2 has multiple ligand binding sites allowing different allosteric modulators to distinctly affect the affinity/efficacy of orthosteric ligands; we recently showed that AZ1729 lacks direct activating effects in its own but turns the natural FFAR2 agonist propionate into a potent activator of the neutrophil NADPH-oxidase, and that neutrophils are activated by the non-activating modulator AZ1729 when combined with another non-activating allosteric modulator, Cmp58 (10). The novel activation/sensitization mediated by the two allosteric FFAR2 modulators was reciprocal, and represented a new regulatory mechanism that controls GPCR signaling. In addition, the down-stream signaling of FFAR2 was biased in that the two interdependent modulators activated the neutrophil NADPH-oxidase, but did not induce a transient rise in the cytosolic concentration of free Ca^2+^([Ca^2+^]_i_); (10)). The results obtained suggested that the two allosteric FFAR2 modulators (Cmp58 and AZ1729) were recognized by FFAR2 through different receptor sites. In order to find new structural variants of allosteric FFAR2 modulators with distinct interaction properties, we selected compounds from the AstraZeneca corporate compound collection with putative FFAR2 interaction profiles and identified ten compounds that were classified as allosteric FFAR2 modulators/agonist. One of these synergistically induced a functionally selective neutrophil activation when combined with AZ1729, but not when combined with Cmp58. The other nine synergistically induced the same type of neutrophil response that was functionally selective but only when combined with Cmp58. No neutrophil activation was achieved when two of the functionally “AZ1729-like” allosteric FFAR2 modulators were combined, supporting the model in which FFAR2 has two sites that selectively bind allosteric modulators. The novel neutrophil activation patterns and receptor down-stream signaling mediated by two cross-sensitizing allosteric FFAR2 modulators represent a new regulatory mechanism that controls receptor function.

## Results

### Neutrophil activation patterns induced by orthosteric agonists and allosteric modulators recognized by FFAR2

#### Activation of the neutrophil superoxide generating NADPH-oxidase by orthosteric FFAR2 agonists and the effect of allosteric modulators

No direct activation of the superoxide generating NADPH-oxidase was induced by the natural orthosteric FFAR2 agonists acetate, propionate or butyrate (shown for propionate in Fig 1A), whereas the more potent agonist Cmp1 (14,15) dose-dependently activated the oxidase to generate superoxide (O_2_^−^; Fig 1A). The response induced by orthosteric FFAR2 agonists is substantially increased in neutrophils sensitized with either of the earlier described allosteric FFAR2 modulators AZ1729 or Cmp58 (9). That is, in neutrophils pre-incubated/sensitized with/by an allosteric modulator, propionate was transferred to a potent activating agonist (Fig 1B). Similarly, the response induced by a non-activating concentration of Cmp1 was greatly increased in the presence of the allosteric modulators AZ1729 or Cmp58 (Fig 1C).

**Figure 1.**
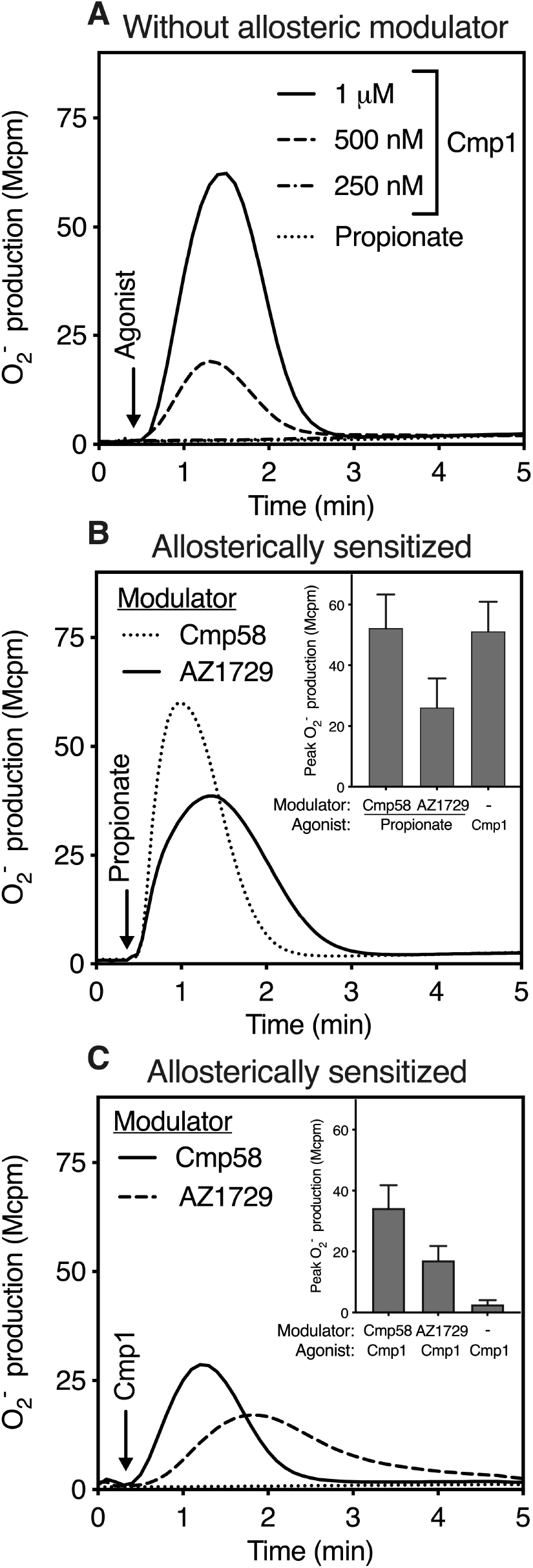
Activation of the neutrophil superoxide (O_2_^−^) generating NADPH-oxidase by orthosteric FFAR2 agonists and effects of the allosteric FFAR2 modulators Cmp58 and AZ1729. **(A)** Human neutrophils were stimulated with either of the two FFAR2 orthosteric agonist Cmp1 (three different concentrations as indicated) or propionate (25 μM; dotted line) and O_2_^−^ production was measured continuously. One representative experiment out of > 5 is shown and time for addition of the agonists is marked by an arrow. **(B)** Production of O_2_^−^ in neutrophils sensitized either with AZ1729 (1 μM for 5 min; solid line) or Cmp58 (1 μM for 5 min; dotted line) when activated by propionate (25 μM; time point for addition indicated by an arrow). Inset: The quantification of O_2_^−^ production when activated by Cmp1 (1μM) alone and by propionate (25 μM) in AZ1729 or Cmp58 sensitized neutrophils (1 μM), respectively. The response was determined from the peak activity and expressed in mega counts per minute (Mcpm); mean ± SD, n = 3. **(C)** Production of O_2_^−^ in neutrophils sensitized with AZ1729 (1 μM for 5 min; broken line) or Cmp58 (1 μM for 5 min; solid line) when activated with a non-activating concentration of Cmp1 (250 nM). For comparison, the response induced by Cmp1 (250 nM) in neutrophils not sensitized with an allosteric modulator is included (dotted line). Inset: Quantification of the neutrophil activity when activated by Cmp1 (250 nM) in AZ1729 (1 μM), Cmp58 (1 μM) sensitized, or non-sensitized neutrophils, respectively. The response was determined from the peak activity and expressed in Mcpm; mean ± SD, n = 3.

#### Neutrophil activation patterns by two allosteric FFAR2 modulators

The two earlier described allosteric modulators Cmp58 and AZ1729 (9) have no direct NADPH-oxidase activating effects in neutrophils, but as mentioned they both turn propionate into a potent activating agonist (Fig 1). More importantly and in agreement with earlier findings (10), the two allosteric modulators, when combined, trigger an activation of neutrophils and superoxide is generated in the absence of any orthosteric agonist; AZ1729 turned Cmp58 into a potent neutrophil activating ligand (Fig 2A), and this activation was reciprocal as Cmp58 turned AZ1729 into a potent activating ligand (Fig 2B). Taken together, the unique neutrophil activation patterns described for the orthosteric FFAR2 agonists and the two allosteric FFAR2 modulators were used to characterize and classify fifteen compounds included in the study.

**Figure 2.**
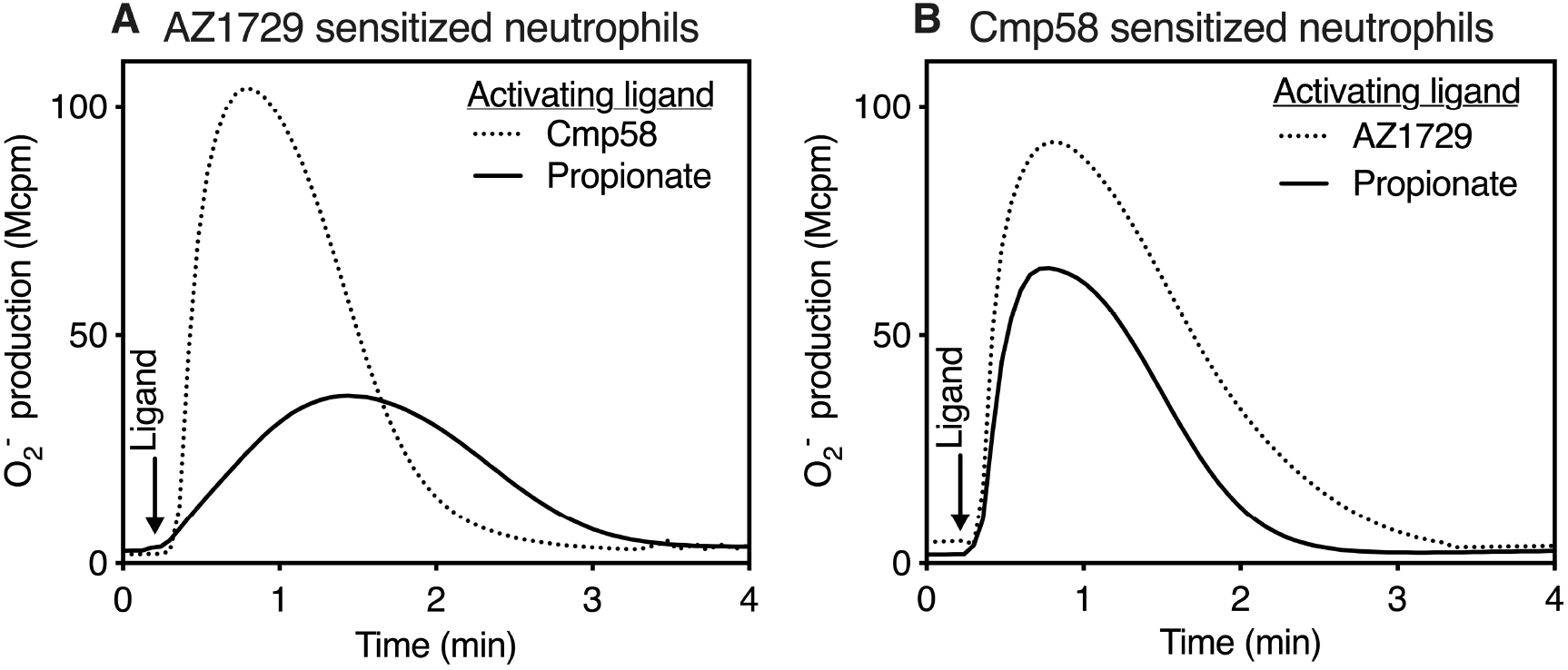
Activation of neutrophils sensitized with the allosteric FFAR2 modulators Cmp58 and AZ1729. **(A)** Human neutrophils were sensitized with the non-activation allosteric FFAR2 modulator AZ1729 (1 μM for 5 minutes) and then activated with either the orthosteric FFAR2 agonist propionate (25 μM; solid line) or the allosteric FFAR2 modulator Cmp58 (1 μM; dotted line) and O_2_^−^ production was measured continuously. One representative experiment out of > 5 is shown and time for addition of the activating ligands is marked by an arrow. **(B)** Human neutrophils were sensitized with the non-activation allosteric FFAR2 modulator Cmp58 (1 μM for 5 minutes) and then activated with either the orthosteric FFAR2 agonist propionate (25 μM; solid line) or the allosteric FFAR2 modulator AZ1729 (1 μM; dotted line) and O_2_^−^ production was measured continuously. One representative experiment out of > 5 is shown and time for addition of the activating ligands is marked by an arrow.

### FFAR2 ligands identified through a neutrophil activation screen of small compounds

#### Direct neutrophil activation

In this study we included: A) Five (AZ4626, AZ0682, AZ0688, AZ1725 and AZ2282) slightly basic thiazole guanidines as close analogs of AZ1729; B) four basic dihydroisoquinoline analogs (AZ8703, AZ1667, AZ1702 and AZ7863), which are structurally close in chemical space but structurally unrelated to both Cmp58 and AZ1729; C) three neutral analogs of Cmp58 (AZ3951,AZ7004 and AZ5994, the latter being identical to Cmp2 and 44, respectively, in (16); AZ3951 and AZ7004 are an enantiomeric pair); D) a group of carboxylic acids of which two (AZ4357 and AZ6732) are structurally related to Cmp1, which in turn share structural elements of Cmp58 and analogs. CATPB and AZ1227 are also carboxylic acids but structurally diverse and also diverse relative all compounds used. Some of the agonists have been described earlier (12,15–20) and all analogs previously not disclosed, were originally identified in FFAR compound screens at AstraZeneca (structures shown in Fig 3A and B). The selected compounds were first screened for their potential to directly activate neutrophils to produce superoxide.

**Figure 3.**
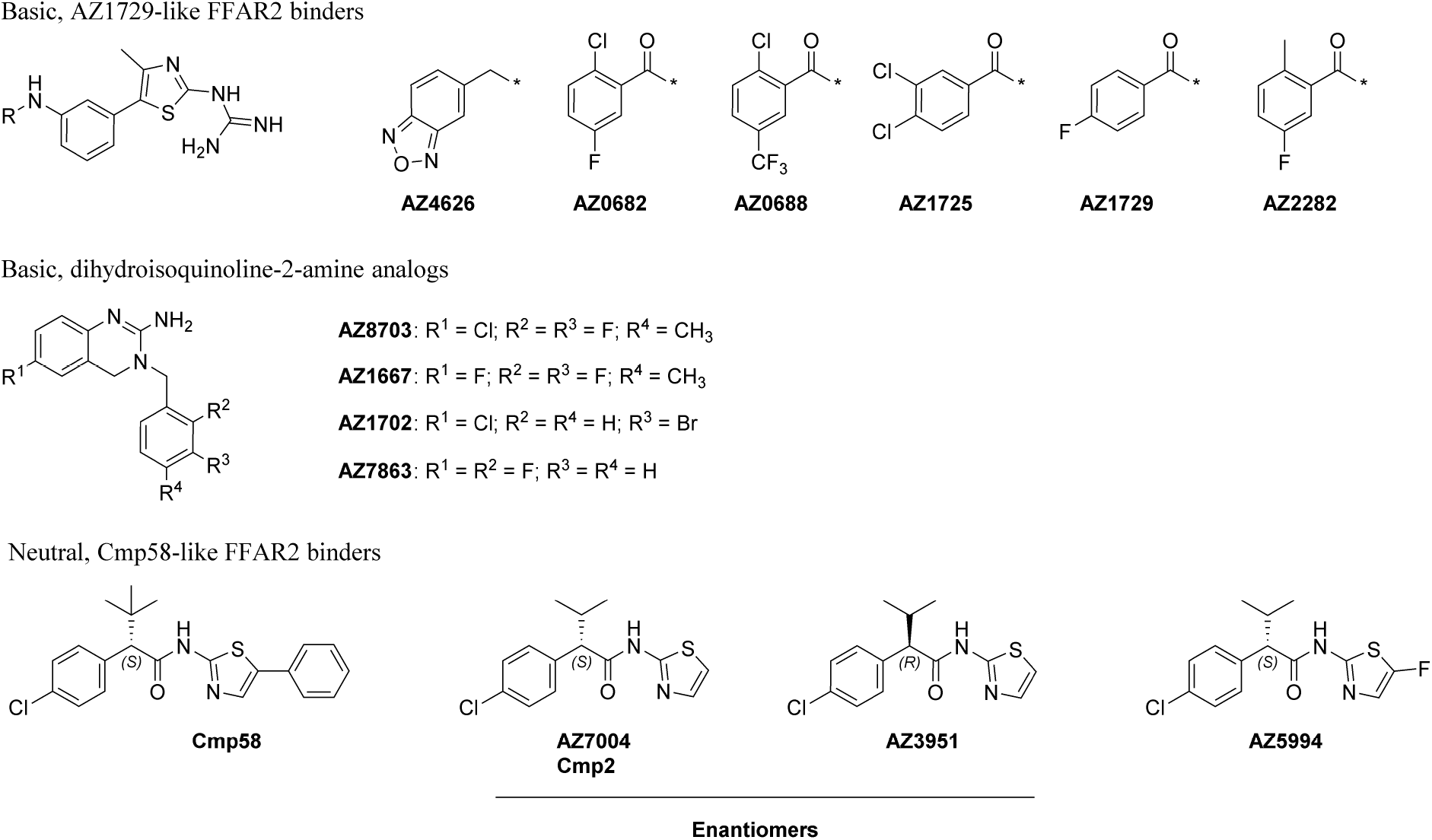
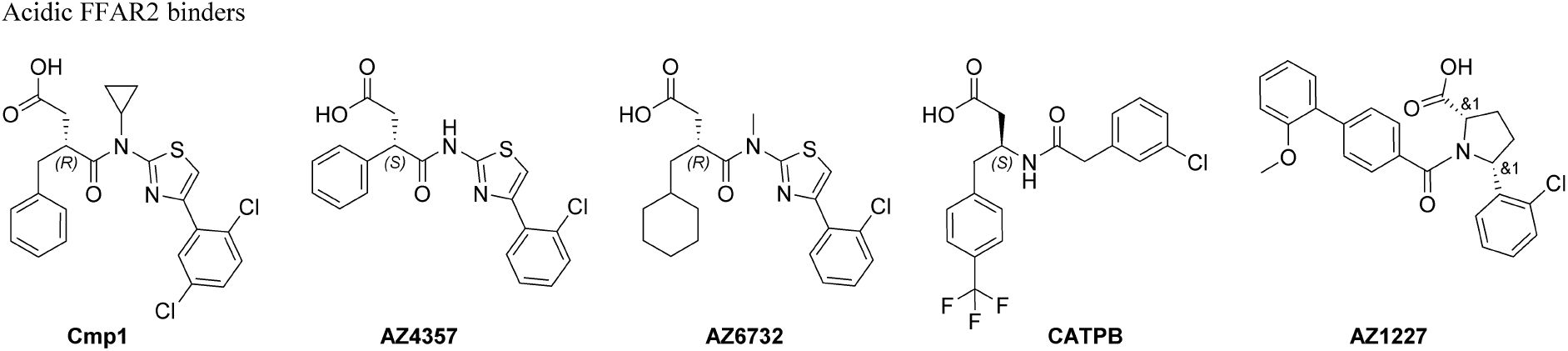
Structure of the compounds used in the study. **(A)** Basic and neutral compounds **(B)** Acidic compounds

It is clear from the data obtained, that although the response induced was fairly low, two (AZ5994 and AZ6732) out of the 15 compounds included in the study activated neutrophils to produce superoxide (Fig 4A). The propionate induced response in neutrophils sensitized with the allosteric FFAR2 modulator Cmp58 was used for comparison as positive control (100%; see Fig 4A, inset). The FFAR2 selective antagonist CATPB reduced the response induced by the two activating compounds (Fig 4B and C), suggesting that they are orthosteric FFAR2 agonists.

**Figure 4.**
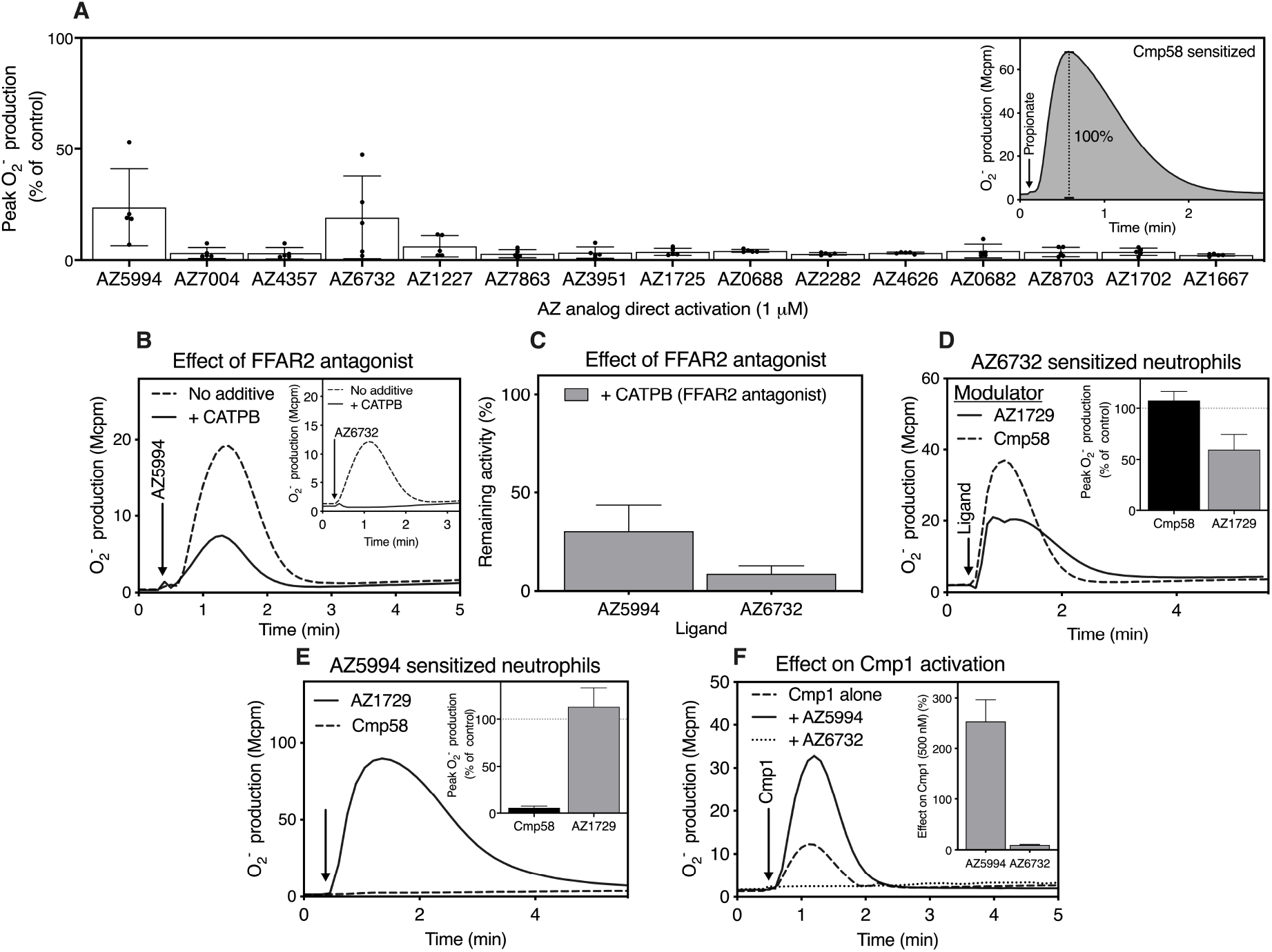
**(A)** *Direct neutrophil activating effects of the presumed allosteric FFAR2 modulators* Non FFAR2 sensitized (naïve) human neutrophils were activated with the different presumed allosteric FFAR2 modulators (1 μM each) and O_2_^−^ production was measured continuously. The peak activities were determined and given in percent of the peak values of the response induced by propionate (25 μM) in neutrophils sensitized (1 μM for 5 min) with Cmp58. Five independent experiments were performed and the results are given as mean ± SD. **Inset:** The response induced by propionate (the value used as 100% reference) in Cmp58 sensitized neutrophils. **(B)** *The response in* naïve *neutrophils induced by AZ5994 and AZ6732 is inhibited by the FFAR2 antagonist CATPB*. Naïve neutrophils pre-incubated without (broken line) or with the FFAR2 antagonist CATPB (100 nM for 5 min; solid line) were activated with AZ5954 (1 μM) and O_2_^−^ production was measured. One representative experiment is shown and time for addition of AZ5994 is marked by an arrow. **Inset:** Naïve neutrophils pre-incubated without (broken line) or with the FFAR2 antagonist CATPB (100 nM for 5 min; solid line) were activated with AZ6732 (1 μM) and O_2_^−^ production was measured. One representative experiment is shown and time for addition of AZ6732 is marked by an arrow. **(C)** Naïve neutrophils incubated without or with the FFAR2 antagonist CATPB (100 nM for 5 min; solid line) were activated with AZ5954 (1 μM) or AZ6732 (1 μM) and O_2_^−^ production was measured. The peak activities were determined and the results are given as the remaining activity (with antagonist in % of without antagonist; mean ± SD, n = 3). **(D)** Neutrophils were sensitized with AZ6732 (1 μM for 5 min) and then activated by AZ1729 (1 μM; solid line) or Cmp58 (1 μM; broken line) and O_2_^−^ production was measured continuously. One representative experiment is shown and the time for addition of AZ1729 or Cmp58 is indicated by an arrow. **Inset:** The response induced in AZ6732 sensitized (1 μM for 5 min) neutrophils when activated by AZ1729 (1μM) and Cmp58 (1μM), respectively. The response was determined from the peak activity and expressed in percent of the activity obtained with Cmp58 sensitized neutrophils activated with AZ1729; mean ± SD, n = 3. **(E)** Neutrophils were sensitized with AZ5994 (1μM for 5 min; solid line) and then activated by AZ1729 (1μM; solid line) or Cmp58 (1 μM; broken line) and O_2_^−^ production was measured continuously. One representative experiment is shown and the time for addition of AZ1729 or Cmp58 is marked by an arrow. **Inset:** The response induced in AZ5994 (1 μM) sensitized (1μM for 5 min) neutrophils when activated by AZ1729 (1 μM) and Cmp58 (1 μM), respectively. The response was determined from the peak activity and expressed in percent of the activity obtained with Cmp58 sensitized neutrophils activated with AZ1729; mean ± SD, n = 3. **(F)** The Cmp1 induced response is enhanced in AZ5994 sensitized neutrophils and reduced in AZ6732 sensitized cells. Neutrophils were sensitized with AZ5994 (1μM for 5 min; solid line) or AZ6732 (1 μM for 5 min; dotted line) and then activated by Cmp1 (500 nM) and O_2_^−^ production was measured continuously. One representative experiment is shown, the time for addition of Cmp1 is marked by an arrow and, for comparison, the response induced by Cmp1 (500 nM; dashed line) in non-sensitized (naïve) neutrophils is shown. **Inset:** The response induced by Cmp1 (500 nM) in neutrophils sensitized with AZ5994 (1 μM) or AZ6732 (1 μM). The response was determined from the peak activity and expressed in percent of the response induced by Cmp1 in naïve neutrophils; mean ± SD, n = 3.

To further study the orthosteric binding mode of AZ5994 and AZ6732, we applied the neutrophil activation pattern described for orthosteric agonists, i.e., the expected outcome for the response induced by pure orthosteric agonists should be substantially increased in neutrophils allosterically modulated by either of the two modulators Cmp58 and AZ1729 (see Fig 1). Indeed, this was also the outcome when neutrophils sensitized with AZ6732 were activated with either of the allosteric modulators AZ1729 or Cmp58 (Fig 4D), confirming that AZ6732 is an orthosteric FFAR2 agonist, which unlike AZ4357 also share the alkylated amide and (*R*)-stereochemistry with the FFAR2 orthosteric agonist Cmp1. Also, neutrophils sensitized with AZ5994 were activated by the allosteric modulator AZ1729, but not when combined with Cmp58 (Fig 4E). This suggests that AZ5994 is not a classical orthosteric agonists but should possibly be classified as an allosteric FFAR2 agonist that, similar to Cmp58, activate neutrophils when combined with AZ1729. The classification of AZ6732 and AZ5994 as an orthosteric agonist and an allosteric agonist, respectively, gained further support from the fact that the neutrophil response induced by Cmp1 was reduced by AZ6732 whereas AZ5994 potentiated this response, data that strengthen the conclusion that AZ5994 is an allosteric FFAR2 modulator with agonistic properties (Fig 4F). It is worth mentioning, that AZ5994 and AZ7004 are structurally a very close analogs and both are relatively close analogs of Cmp58. The 13 compounds that lacked the ability to activate the NADPH-oxidase (Fig 4A), could have been inert with respect to the ability to interact with FFAR2 at all, but there was also the possibility that some were allosteric FFAR2 modulators.

#### Allosteric FFAR2 modulators identified through a positive (sensitizing) effect on the neutrophil response induced by the orthosteric agonist propionate

All 15 compounds were used as neutrophil sensitizers and after a 5 min pre-incubation with respective compound, the neutrophils were activated with propionate and the ability of the neutrophils to produce superoxide was determined. The response induced by propionate in AZ1729 modulated neutrophils was used as positive control for comparison (Fig 5, inset). In accordance with the data presented above, AZ5994, the compound suggested to be an allosteric agonist, transferred propionate to a neutrophil activating agonist (Fig 5). Based on the activation profile of AZ5994 (Fig 4), this was an effect that was expected and also in agreement with the suggestion that this is an allosteric FFAR2 agonist also having modulating effects. The fact that the direct activating compound AZ6732 belonged to the group of compounds that lacked the ability to change propionate into an oxidase activating agonist (Fig 5), further strengthens the conclusion that AZ6732 is not an allosteric modulator but rather an orthosteric agonist.

**Figure 5.**
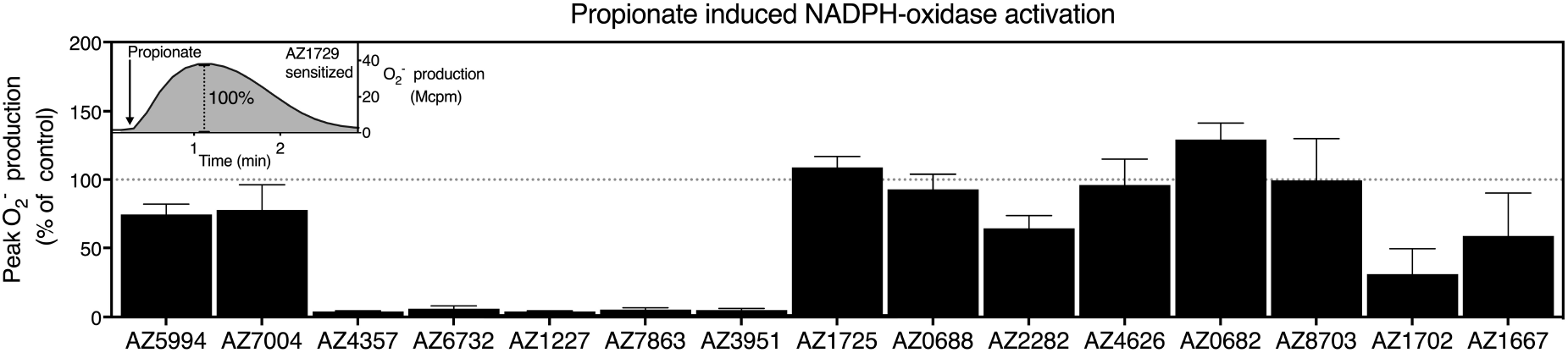
The response induced by propionate in neutrophils sensitized with the presumed allosteric FFAR2 modulators. Human neutrophils sensitized with the presumed allosteric FFAR2 AZ-modulators (1 μM each, 5 min) were activated by propionate (25 μM) and O_2_^−^ production was measured continuously. The peak activities were determined and expressed in percent of the peak value of the propionate induced response in neutrophils sensitized (1μM for 5 min) with the allosteric FFAR2 modulator Cmp58. Three independent experiments were performed and the results are given as men ± SD. **Inset:** The response induced by propionate (the value used as 100% reference) in Cmp58 sensitized neutrophils.

### Characterization of the synergistic activation pattern of new allosteric FFAR2 modulators

#### Neutrophil sensitization by non-direct activating compounds also lacking effect on the propionate response

The four non-direct activating compounds that also lacked the ability to increase the activity induced by propionate (AZ4357; AZ1227; AZ3951; AZ7863) could still be weak orthosteric FFAR2 agonists, and in order to investigate this possibility the effects of the allosteric modulators (Cmp58 and AZ1729) were determined. Neutrophils were sensitized with respective compound and then activated with either Cmp58 (Fig 6A) or AZ1729 (Fig 6B) and the ability of the neutrophils to produce superoxide was determined. According to the neutrophil activation pattern described above, Cmp58 as well as AZ1729 should activate neutrophils treated/activated with an orthosteric agonists, and it is clear from the data presented that no activity was induced with AZ3951 using either of the allosteric modulators, suggesting that this compound lacks the ability to interact with FFAR2 (Fig 6A, B). It should also be noted that AZ3951 is the (*R*)-enantiomer of AZ7004 and confirms the importance of stereochemistry on the Cmp58 series to bind FFAR2. In contrast, the NADPH-oxidation activation induced both by Cmp58 and AZ1729 were increased in AZ4357 and AZ1227 sensitized neutrophils, suggesting that the latter two compounds are orthosteric agonists. Although the activity is low, the response induced by Cmp58 was increased in AZ7863 sensitized neutrophils (Fig 6A); no such activating effect was obtained with AZ1729 (Fig 6B), suggesting that this compound should be classified as a weak allosteric modulator that is functionally”AZ1729-like”.

**Figure 6.**
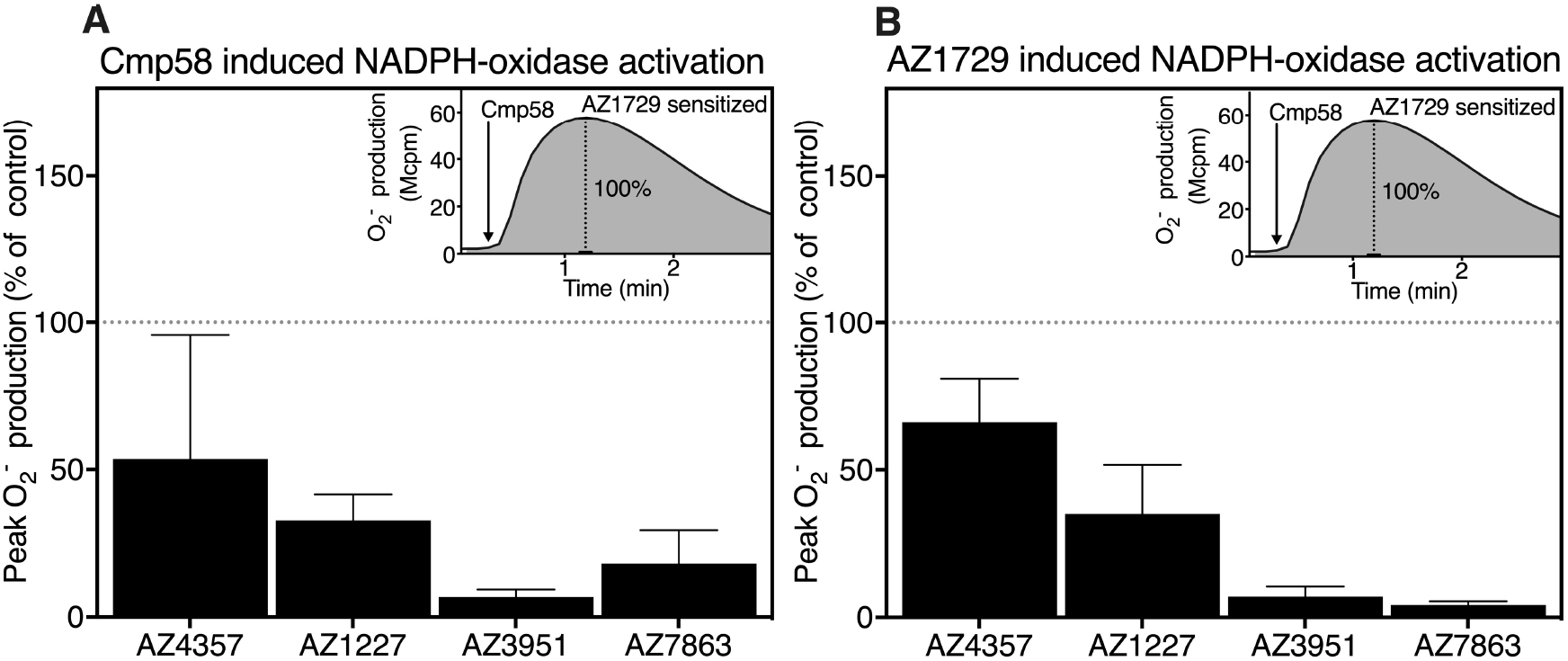
Response induced by Cmp58 and AZ1729 in neutrophils sensitized with the compounds being unable to modulate the propionate response. (**A and B**) Neutrophils sensitized with the compounds (1 μM each) found to be unable to modulate the propionate response (see Fig 5) were activated by Cmp58 (1 μM, **A**) or AZ1729 (1 μM, **B**), and O_2_^−^ production was measured continuously. The peak activities were determined and expressed in percent of the peak value of the Cmp58 (1 μM) induced response in neutrophils sensitized with the allosteric FFAR2 modulator AZ1729 (1 μM for 5 min). Three independent experiments were performed and the results are given as men ± SD. **Inset in A and B:** The response induced by Cmp58 (the value used as 100% reference) in AZ1729 sensitized neutrophils.

#### Neutrophil activation induced by Cmp58

When the two different allosteric binding sites in FFAR2 (the site for Cmp58 and that for AZ1729) are occupied simultaneously, neutrophils are activated to produce superoxide (Fig 2). To determine the binding site for the compounds shown to be allosteric modulators, a classification based on the sensitizing effect on the propionate induced response (see Fig 5), the activation patters were determined using Cmp58 as the neutrophil activating ligands. The new allosteric modulating compounds were used to sensitize neutrophils and following a 5 min incubation period the cells were activated with Cmp58 (Fig 7A) and the ability of the neutrophils to produce superoxide was determined. According to the neutrophil activation pattern described above, Cmp58 should activate neutrophils allosterically modulated with compounds that interact with the same binding site as AZ1729. It is clear from the data presented that AZ5994 and AZ7004 were without effect whereas neutrophils sensitized with the other 8 compounds were activated by Cmp58 (Fig 7A). These data suggest that AZ5994 and AZ7004 are functionally “Cmp58-like”.

**Figure 7.**
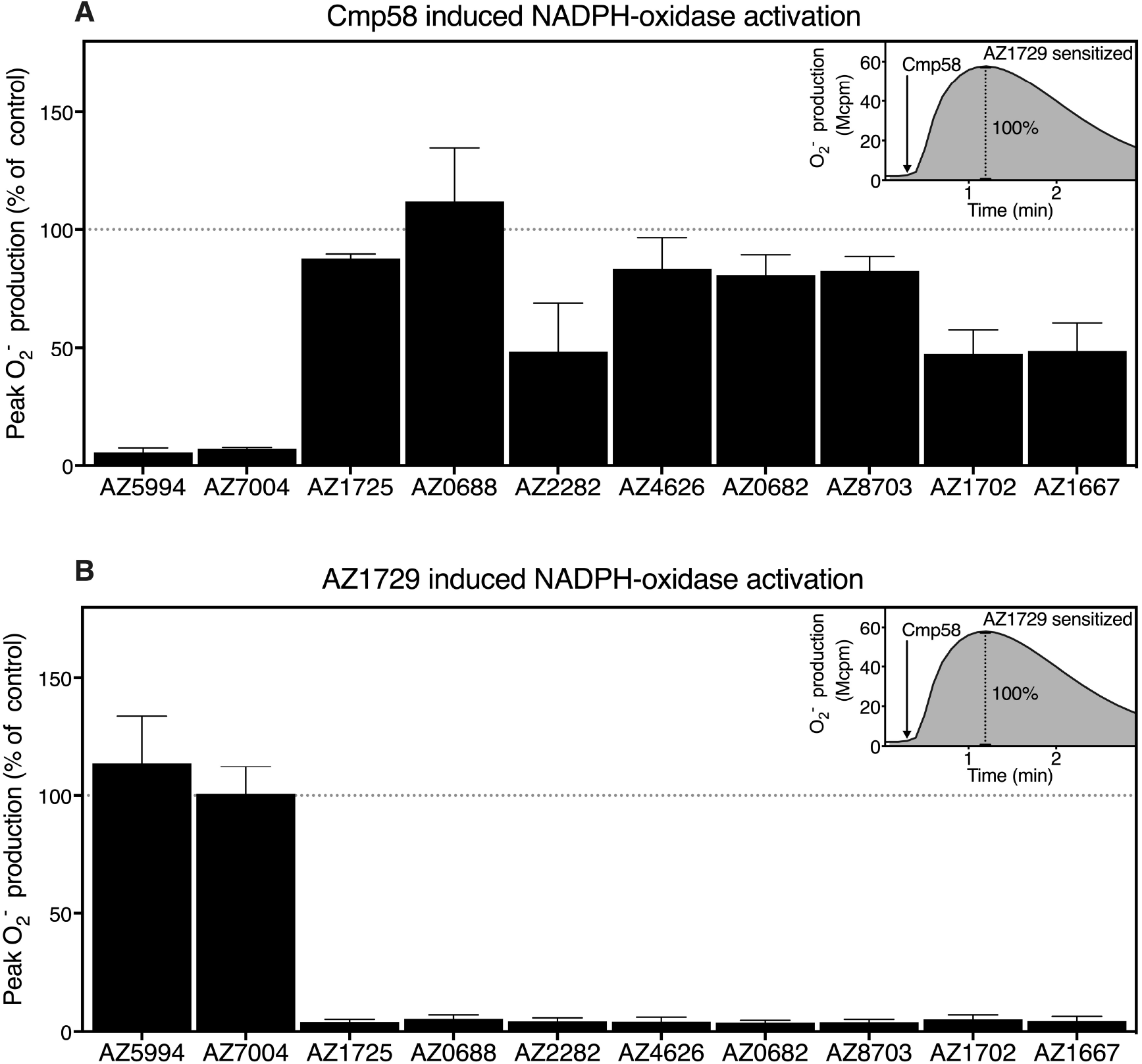
Classification of the compounds found to modulate the neutrophil response induced by propionate. **(A)** Neutrophils sensitized with the with the compounds (1 μM each) found to positively modulate the neutrophil response induce by propionate (see Fig 5) were activated by Cmp58 (1 μM) and O_2_^−^ production was measured continuously. The peak activities were determined and expressed in percent of the peak value of the Cmp58 (1 μM) induced response neutrophils sensitized with the allosteric FFAR2 modulator AZ1729 (1 μM for 5 min). Three independent experiments were performed and the results are given as men ± SD. **Inset:** The response induced by Cmp58 (the value used as 100% reference) in AZ1729 sensitized neutrophils. **(B)** Neutrophils sensitized with the with the compounds (1 μM each) found to positively modulate the neutrophil response induce by propionate (see Fig 5) were activated by AZ1729 (1 μM) and O_2_^−^ production was measured continuously. The peak activities were determined and expressed in percent of the peak value of the Cmp58 (1 μM) induced response neutrophils sensitized with the allosteric FFAR2 modulator AZ1729 (1 μM for 5 min). Three independent experiments were performed and the results are given as men ± SD. **Inset:** The response induced by Cmp58 (the value used as 100% reference) in AZ1729 sensitized neutrophils.

#### Neutrophil activation induced by AZ1729

Neutrophils allosterically modulated with a compound that interacts with the same binding site as Cmp58, should in contrast not be activated by Cmp58, but instead be activated by AZ1729. It is clear from the data presented that neutrophils sensitized with AZ5994 or AZ7004 were selectively activated by AZ1729 (Fig 7B), confirming the suggestion that these two allosteric modulators are functionally “Cmp58-like”. It is also clear from the data presented that the 8 allosteric modulators that were activated by Cmp58 (Fig 7A) were non-active when combined with AZ1729 (Fig 7B), suggesting that these compounds should be classified as allosteric modulators that are functionally “AZ1729-like”.

#### No synergistic activation is achieved with allosteric FFAR2 modulators suggested to interact with the same binding site

According to the model above describing the different neutrophil activation patterns, the two modulators (AZ5994 and AZ7004) that transfer propionate and AZ1729, but not Cmp58, to neutrophil NADPH-activating agonists, are expected to interact with the same binding site as Cmp58. The suggested model gains further support from the fact that no synergistic neutrophil activation was induced when AZ5994 and AZ7004 were combined (Fig 8A), whereas both these compounds sensitized neutrophils when activated with AZ0688 and AZ1702, respectively (Fig 8A), two structurally diverse compounds classified as functionally”AZ1729-like” allosteric modulators.

**Figure 8.**
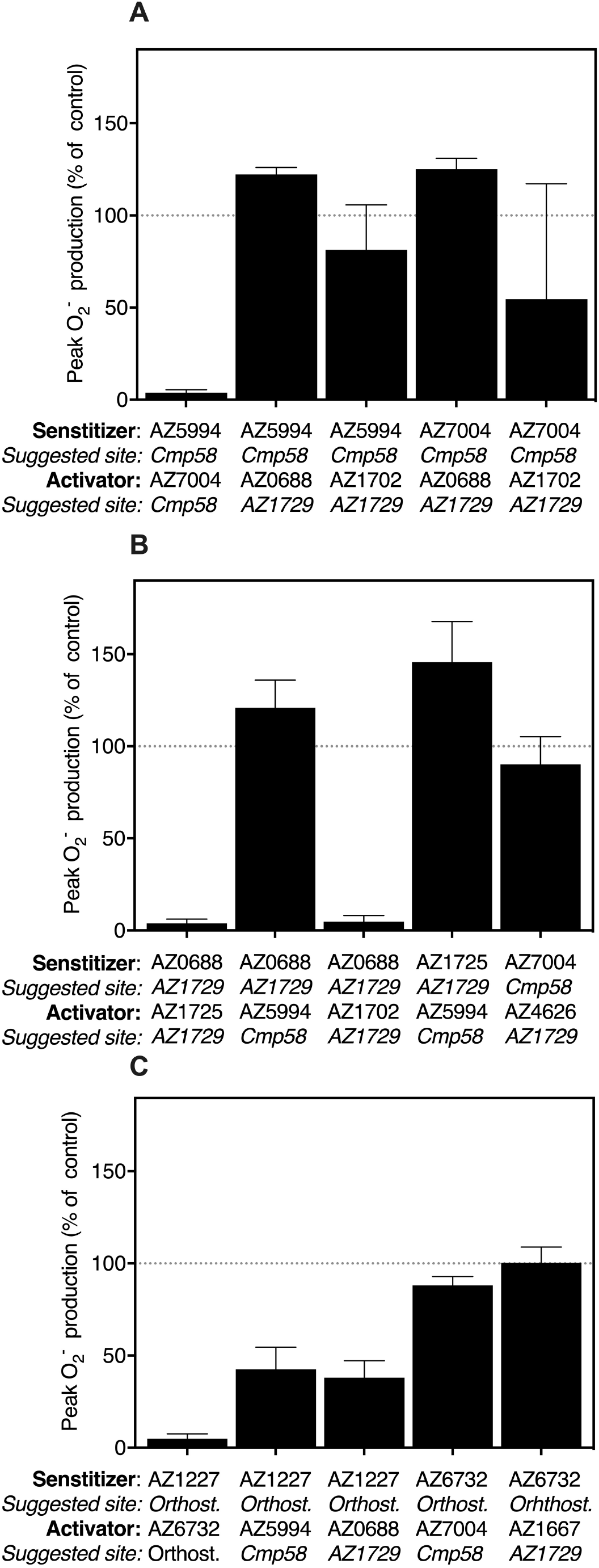
Interdependent neutrophil activation by functionally “Cmp58-like” and “AZ1729-like” compounds. **(A)** Human neutrophils sensitized with a functionally “Cmp58-like” allosteric FFAR2 modulator (1 μM each) were activated by another functionally “Cmp58-like” or a functionally “AZ1729-like” allosteric modulator (1 μM each) and O_2_^−^ production was measured continuously. The presumed binding sites for each sensitizer and activator are indicated. The peak activities were determined and expressed in percent of the peak value of the Cmp58 (1 μM) induced response neutrophils sensitized with the allosteric FFAR2 modulator AZ1729 (1 μM for 5 min). Three independent experiments were performed and the results are given as mean ± SD. **(B)** Human neutrophils sensitized with a functionally “AZ1729-like” allosteric FFAR2 modulator (1μM each) were activated by another functionally “AZ1729-like” or a functionally “Cmp58-like” allosteric modulator (1μM each) and O_2_^−^ production was measured continuously. The presumed binding sites for each sensitizer and activator are indicated. The peak activities were determined and expressed in percent of the peak value of the Cmp58 (1 μM) induced response neutrophils sensitized with the allosteric FFAR2 modulator AZ1729 (1 μM for 5 min). Three independent experiments were performed and the results are given as mean ± SD. **(C)** Human neutrophils sensitized with a presumed “orthosteric-like” allosteric FFAR2 modulator (1μM each) were activated by another orthosteric agonist, a functionally “AZ1729-like” or a functionally “Cmp58-like” allosteric modulator (1 μM each) and O_2_^−^ production was measured continuously. The peak activities were determined and expressed in percent of the peak value of the Cmp58 (1 μM) induced response neutrophils sensitized with the allosteric FFAR2 modulator AZ1729 (1μM for 5 min). Three independent experiments were performed and the results are given as mean ± SD.

It is also clear that no neutrophil activation was induced when one of the allosteric FFAR2 modulators suggested to interact to the same binding site as AZ1729, was combined with another of the new allosteric modulator suggested to bind to the same site (Fig 8B), whereas they activated neutrophils when combined with the functionally “Cmp58-like” compounds AZ5994 and AZ7004, respectively (Fig 8B). In addition, no activation was achieved when a compound suggested to interact with the orthosteric site was combined with another presumed orthosteric agonist, whereas neutrophils were activated when such a compound was combined with a compound presumed to be an allosteric modulator (Fig 8C).

### Signaling biased neutrophil activation by two complementary non-activating allosteric FFAR2 modulators

#### The allosteric modulators lack direct effect but modulates the transient rise in [Ca^2+^]_i_ induced by propionate

Neutrophil activation, measured as a change in [Ca^2+^]_i_, was used to determine the signaling properties of the different FFAR2 modulators (Fig 9). Propionate in high concentrations (250 μM) triggered an activation of the Ca^2+^ signaling pathway in naïve neutrophils, whereas no such effect was seen when the concentration of propionate was reduced to 25 μM (Fig 9A). No direct change in the [Ca^2+^]_i_, was induced by Cmp58 or AZ1729 when used alone to activate neutrophils, but the threshold for the ability of propionate to induce a response was lowered in the presence of either Cmp58 and AZ1729 (Fig 9A; (9)). No direct neutrophil activating effect was induced by any of the allosteric modulators classified as “AZ1729-like” when given alone to naïve neutrophils, but they all lowered the thresh-hold for the propionate response (Fig 9B). A similar pattern was obtained with the FFAR2 ligands classified as “Cmp58-like” with a clean allosteric effect (AZ7004) and an allosteric/agonistic effect (AZ5994), respectively, that is no rise in [Ca^2+^]_i_ was induced by these ligands alone while the threshold for the concentration of propionate was reduced (Fig 9C).

**Figure 9.**
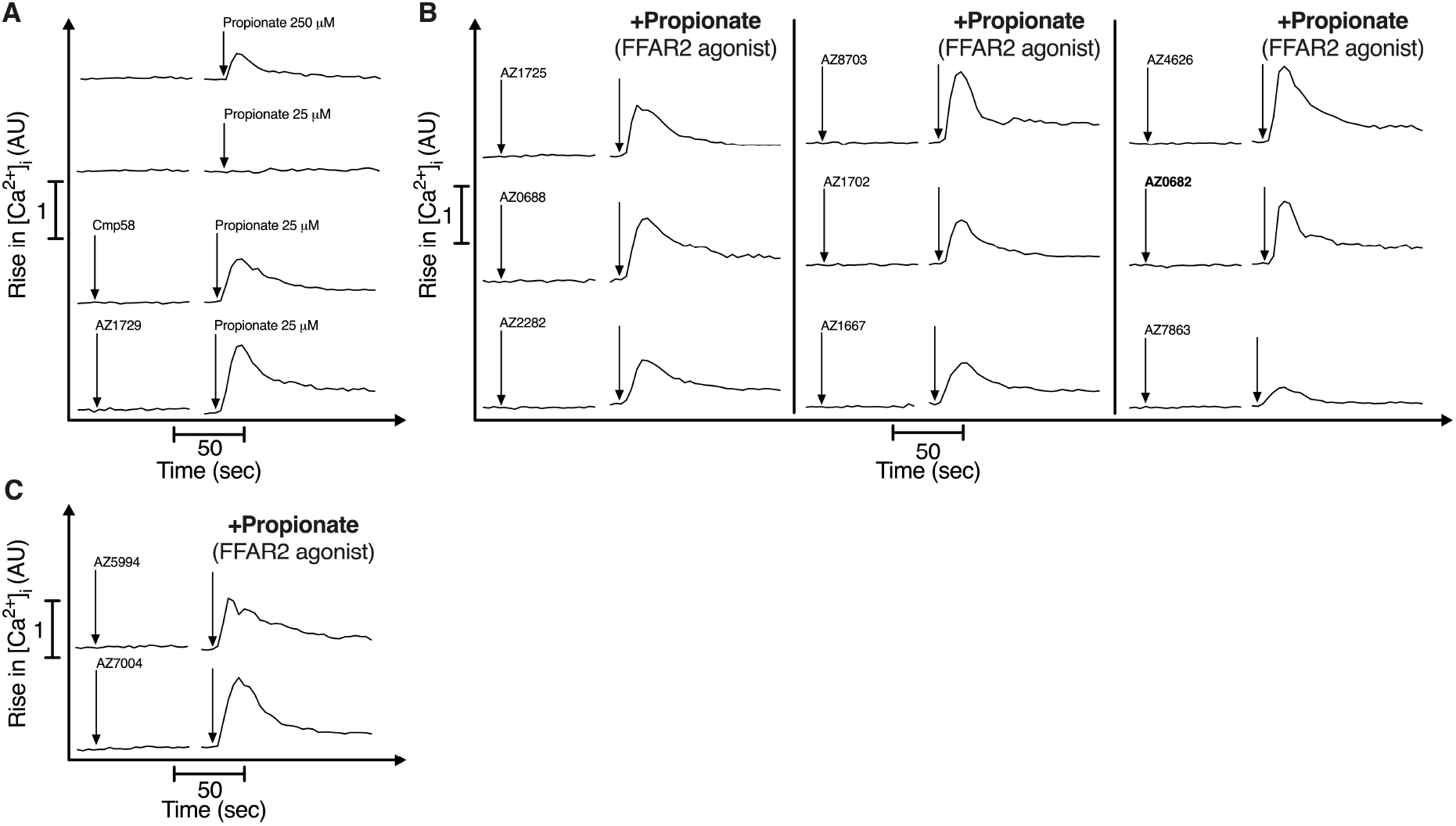
Effects of allosteric modulators on the intracellular concentration of Ca^2+^([Ca^2+^]_i_) in neutrophils. (**A**) The change in [Ca^2+^]_i_ was followed in neutrophils activated either by propionate alone (250 μM and 25 μM; upper part) or by propionate (25 μM) in neutrophil first challenged by Cmp58 (1 μM; lower panel) or AZ1729 (1 μM; lower panel), and the rise in [Ca^2+^]_i_. The time points for addition of the different compounds are indicated by arrows. (**B**) The change in [Ca^2+^]_i_ induced by propionate (25 μM) was followed in neutrophils first challenged by AZ compounds (1 μM) classified as functional “AZ1729-like” (see Fig 7A) The time points for addition of the different compounds are indicated by arrows. (**C**) The change in [Ca^2+^]_i_ induced by propionate (25 μM) was followed in neutrophils first challenged by AZ compounds (1 μM) classified as functional “Cmp58-like” (see Fig 7B) The time points for addition of the different compounds are indicated by arrows.

#### Induction of a transient rise in [Ca^2+^]_i_ when the allosteric modulators are combined

The data presented show that AZ1729 and Cmp58 when added in sequence, activate neutrophils to produce O_2_^−^ (Fig 2), but this activation was achieved without any concomitant rise in [Ca^2+^]_i_ (Fig 10A) and this pattern was valid also when the new allosteric modulators classified as functionally “AZ1729-like” were combined with Cmp58 (Fig 10A), AZ0688 being an exception – a rise in [Ca^2+^]_i_ was induced when AZ0688 was combined with Cmp58 (Fig 10A). Despite the fact that no rise in [Ca^2+^]_i_ was induced by the allosteric agonist/modulator AZ5994 alone, a rise in [Ca^2+^]_i_ was induced when this functionally “Cmp58-like” modulator was combined with AZ1729 (Fig 10B), and such a response was induced also when the second allosteric modulator (AZ7004), classified as functionally “Cmp58-like”, was combined with AZ1729 (Fig 10B). No rise in [Ca^2+^]_i_ was induced by AZ5994 or AZ7004 when Cmp58 replaced AZ1729 as sensitizing modulators, and this was true also for AZ0688 when Cmp58 was replaced by AZ1729 (Fig 10C).

**Figure 10.**
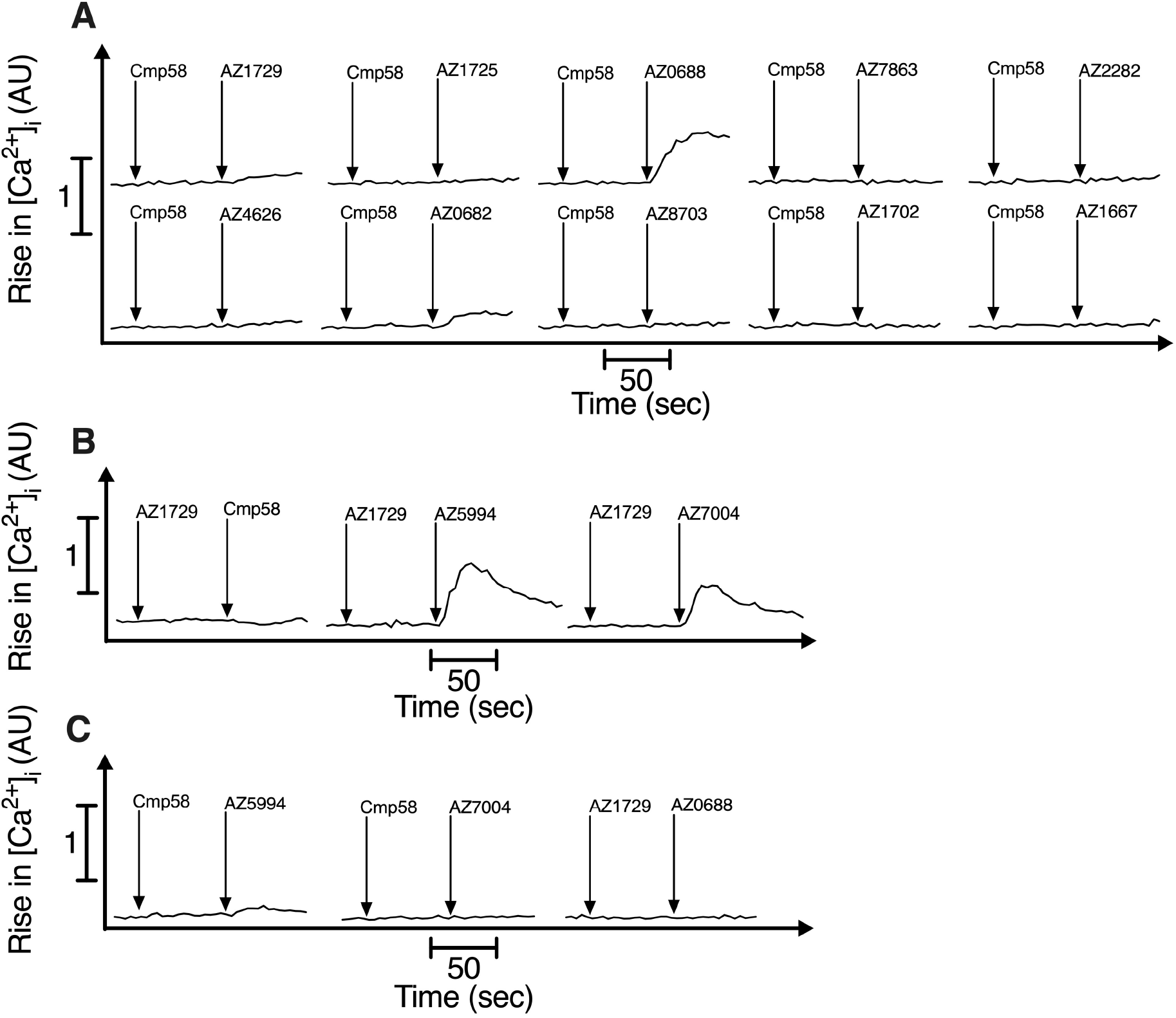
Combined effect of two allosteric modulators on the intracellular concentration of Ca^2+^([Ca^2+^]_i_) in neutrophils. **(A)** The change in [Ca^2+^]_i_ induced by AZ1729 (1μM) and functional “AZ1729-like” allosteric modulators was followed in neutrophils first challenged by Cmp58 (1 μM). The time points for addition of the different compounds are indicated by arrows. **(B)** The change in [Ca^2+^]_i_ induced by Cmp58 (1 μM) and functional “Cmp58-like” allosteric modulators was followed in neutrophils first challenged by AZ1729 (1 μM). The time points for addition of the different compounds are indicated by arrows. **(C)** The change in [Ca^2+^]_i_ induced i) by functional “Cmp58-like” compounds (1μM; AZ5995;AZ7004) was followed in neutrophils first challenged by Cmp58 (1 μM) and ii) by afunctional “AZ1927-like” compounds (1μM; AZ0688) was followed in neutrophils first challenged by AZ1729 (1 μM). The time points for addition of the different compounds are indicated by arrows.

## Discussion

A tentative binding/signaling/activation-model, postulating three receptor binding sites in FFAR2 is presented. This model is based on data showing that the non-activating allosteric FFAR2 modulators AZ1729 and Cmp58 interdependently activate neutrophils to produce/release superoxide (O_2_^-^), and that the signaling profile downstream of the receptor is biased (10) in that no transient rise in intracellular Ca^2+^ is triggered during this activation of FFAR2. Neutrophil activation by orthosteric FFAR2 agonists is modulated in the same way by the non-activating compounds Cmp58 and AZ1729 but the enhanced response is achieved through binding of the two modulators to distinctly different allosteric receptor sites. The precise FFAR2 sites recognizing Cmp58 and AZ1729, respectively, are not known. However, the experimental data in this study, obtained through characterization of different FFAR2 ligands strengthen the model surmising three distinct binding sites. Signaling by FFAR2 is complex, illustrated by the fact that when orthosteric agonists are involved in activation, the signaling generated downstream of the receptor activates the neutrophil oxidase as well as the Ca^2+^pathway, whereas signaling without a direct involvement of the orthosteric binding site is biased away from the Ca^2+^ pathway. It is also worth noticing, that the FFAR2 ligands AZ5994 and AZ7004 used in this study were found to be allosteric FFAR2 agonists having also an allosteric modulating effect similar to that of Cmp58.

By definition, the non-direct-activating molecules included in the study that positively modulate the neutrophil response induced by the orthosteric FFAR2 agonist propionate, should be classified as allosteric FFAR2 modulators. And according to our binding site model, these allosteric modulators should be expected to activate FFAR2 when combined with one but not the other of the earlier described allosteric FFAR2 modulators Cmp58 and AZ1729. In line with the model, allosteric FFAR2 modulators that synergistically induce a signaling biased neutrophil activation when combined with Cmp58, were non-activating when they were combined with AZ1729. In addition, and also in line with the model, the activation signaling down-stream of FFAR2, when the receptor was activated by Cmp58 together with a ligand found to be functionally “AZ1929-like”, was biased. That is, the two complementary modulators together triggered an activation of the NADPH-oxidase but not any transient rise in the cytosolic concentration of [Ca^2+^]_i_. No neutrophil activation was induced when two of the functionally “AZ1729-like” allosteric FFAR2 modulators were combined. Taken together all these data support the FFAR2 activation model in which the receptor has two sites that selectively recognize different allosteric modulators. The allosteric modulators classified as functionally “AZ1729-like” belong to two different chemical series, one represented by structures similar to AZ1729 while the other structurally diverse series consists of dihydroisoquinolines and represent a new series of FFAR2 ligands (Table 1 and Fig 11). It is noteworthy that these diverse chemical series interact with the same allosteric binding site on the receptor and there were no functional differences between the compound groups. It should be noticed that although AZ7863 is a close analog to AZ8703, AZ1702 and AZ1667, it lacks all effects on FFR2, which suggests the meta substituent (R3, Fig 3A) should not be a hydrogen for an interaction with FFAR2, but this remains to be proven.

**Table 1.** Basic characteristics and classification of the compounds included in the study.

| Compound | Modulation of |  |  |  | Classification |
| --- | --- | --- | --- | --- | --- |
|  | Neutrophil activation | Propionate | Cmp58 | AZ1729 |  |
| **Cmp1** | + | - | + | + | orthost. agonist |
| **Cmp58** | - | + | - | + | Cmp58 AM |
| **AZ1729** | - | + | + | - | AZ1729 AM |
| **CATPB** | - | - | - | - | orthost. antagonist |
| **AZ5994** | + | + | - | + | Cmp58-like agonist |
| **AZ7004** | - | + | - | + | Cmp58-like AM |
| **AZ4357** | - | - | + | + | orthost. agonist |
| **AZ7863** | - | - | (+) | - | AZ1729-like |
| **AZ6732** | + | - | + | + | orthost. agonist |
| **AZ1227** | - | - | + | + | orthost. agonist |
| **AZ3951** | - | - | - | - | not recognized |
| **AZ2282** | - | + | + | - | AZ1729-like AM |
| **AZ1725** | - | + | + | - | AZ1729-like AM |
| **AZ0688** | - | + | + | - | AZ1729-like AM |
| **AZ4626** | - | + | + | - | AZ1729-like AM |
| **AZ0682** | - | + | + | - | AZ1729-like AM |
| **AZ8703** | - | + | + | - | AZ1729-like AM |
| **AZ1702** | - | + | + | - | AZ1729-like AM |
| **AZ1667** | - | + | + | - | AZ1729-like AM |

AM = allosteric modulator; “+” indicates neutrophil superoxide production; “-” denotes no response.

**Figure 11.**
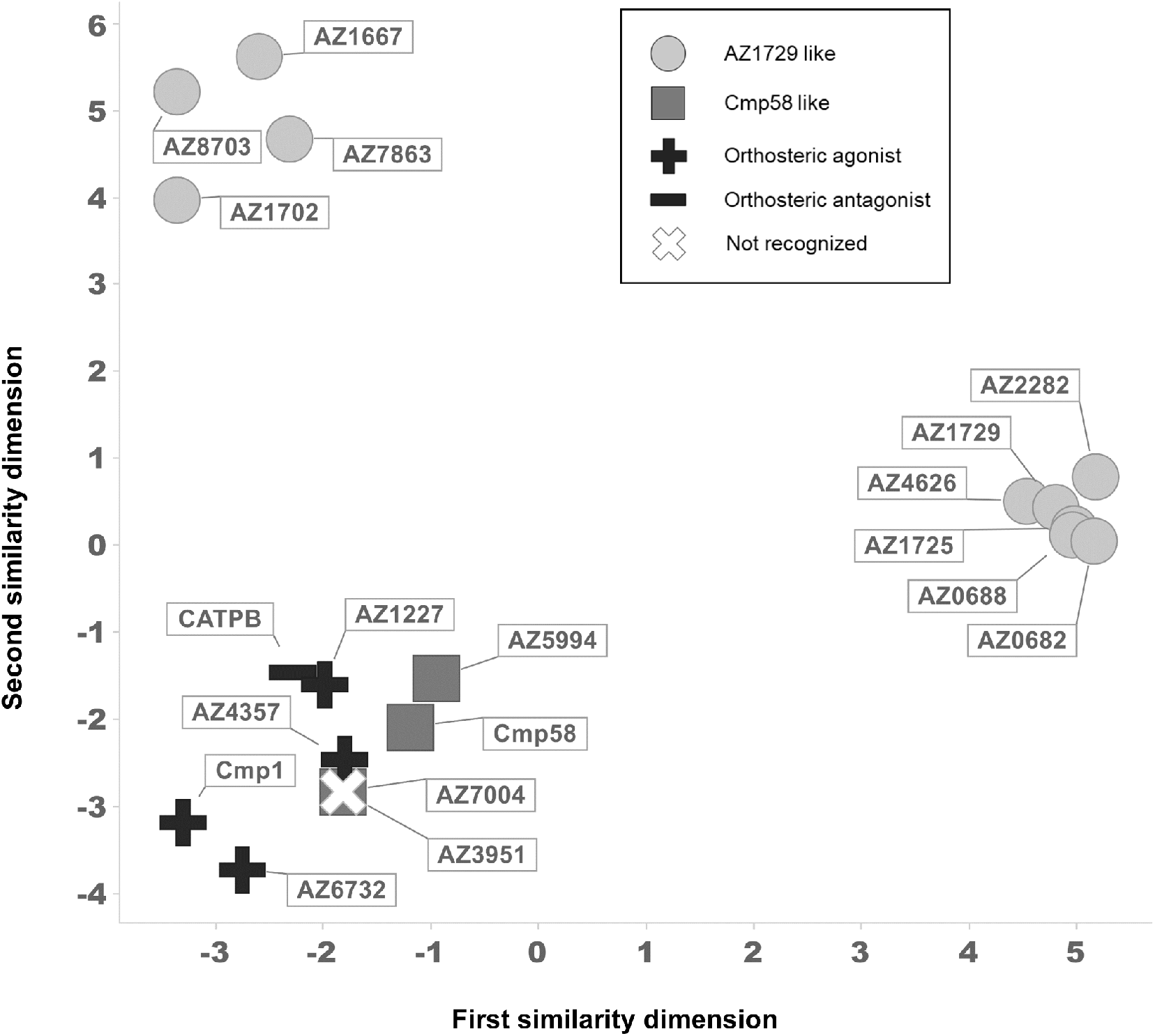
A two-dimensional structural similarity map between the compounds included in the study and the suggested FFAR2 classification. Extended-connectivity fingerprints (ECFP_6) were used to calculate Tanimoto scores and generate a two-dimensional structural similarity map. Structures close in both dimensions share structural features. Shape and color represent the classifications used in Table 1.

According to the three binding site model, a functionally “Cmp58-like” allosteric FFAR2 modulator should induce the same response pattern as that described above for Cmp58 when this compound is combined with an orthosteric FFAR2 agonist such as propionate and with AZ1729. The molecule AZ7004 used in the study, is the *S*-enantiomer of an earlier described FFAR2 ligand, the phenylacetamide 4CMTB (also named AMG7703). AZ7004 was shown earlier to be an FFAR2 agonists that also enhances the ability of acetate to induce a rise in [Ca^2+^]_i_ (20); the compound has thus been classified as an allosteric FFAR2 agonist. Even if no direct neutrophil activating effects was induced by AZ7004, the results obtained are in agreement with the earlier suggested classification of 4CMTB. It is clear from the results obtained, that AZ7004 positively modulates the response induced by propionate, and an agonistic effect of AZ7004 is observed when combined with AZ1729 as illustrated not only by the AZ1729 mediated sensitization of the oxidase activity induced by AZ7004, but also by the [Ca^2+^]_i_ rise induced by AZ7004 in AZ1729 sensitized neutrophils. It is clear that Cmp58 had no modulating effect on the response induced by AZ7004, suggesting that this compound should be classified i) as a functionally “Cmp58-like” allosteric modulator, and ii) as an agonist with binding/activation properties that differ from the classical orthosteric agonists. The largest structural difference between Cmp58 and AZ7004 is a phenyl versus hydrogen in the 5-position of the thiazolyl group. While this does not impact the ability to positively modulate the response induced by propionate or AZ1729 in oxidase activity, the ability to function as an agonist is only added in the absence of the 5-phenyl and this is shown as an ability to trigger a transient rise in [Ca^2+^]_i_ in AZ1729 sensitized neutrophils. The fact that AZ3951, the (*R*)-enantiomer of AZ7004 ((*R*)-4CMTB) lacks both activating and modulating effects on FFAR2 is in agreement with earlier findings showing that (*R*)-4CMTB is less potent that (*S*)-4CMTB (21).

Cmp58 as well as AZ7004 and AZ5994 were originally profiled in the same screen designed to identify small molecule FFAR2 ligands (16), and structurally AZ5994 is very similar to AZ7004 (F versus H respectively, Fig 3) and consequently AZ5994 should be classified an allosteric FFAR2 agonist. Despite the fact that AZ5994, in contrast to AZ7004, activates the neutrophil NADPH-oxidase, the basic functional properties of these two compounds are the same. It is clear from the results obtained, that AZ5994 enhances (positively modulates) the response induced by both propionate and Cmp1, and has an agonistic effect when combined with AZ1729; this is evident as a transient rise in [Ca^2+^]_i_ induced in AZ1729 sensitized neutrophils. Cmp58 had no modulating effect on the response induced by AZ5994, and taken together, also AZ5994 should be classified as a “Cmp58-like” allosteric modulator. The fact that the direct activating effect was only partly inhibited by the FFAR2 antagonist CATPB and that the response induced by orthosteric FFAR2 agonists (propionate and Cmp1) was positively modulated, suggest that the agonistic property (recognition/binding) of AZ5994 differs from that of classical orthosteric agonists.

The results presented show that FFAR2 can adopt two different conformational states with different signaling properties; i) the signals generated by allosterically modulated FFAR2s, when activated by an othosteric agonist, trigger an assembly of the neutrophil NADPH-oxidase as well as those that activate the PLC-PIP_2_-IP_3_-Ca^2+^ pathway, whereas, ii) the signals generated by allosterically modulated receptors, when activated by a second allosteric modulator, recognized by a different receptor site, may trigger an assembly of the oxidase but not those that activate the PLC-PIP_2_-IP_3_-Ca^2+^ pathway, and, even though the precise signals that initiate an assembly of the NADPH-oxidase components and an activation of the system are unknown, it is thus clear that the PLC-PIP_2_-IP_3_-Ca^2+^ pathway is not needed for an activation of the oxidase. The tool allosteric modulators AZ1729 and Cmp58 used in the study were both initially identified in screening studies designed to identify FFAR2 ligands (12,16), and the fact that they both lack the carboxylic acid functionality needed for orthosteric agonists to activate FFAR2 (22), implies that they interact with allosteric receptor sites. This is in agreement with the fact both these modulators as well as all the “AZ1729 analogs exerted similar modulating functions and also lacked detectable direct neutrophil activating effects characteristic for an orthosteric agonists. The acidic group is missing also in the Cmp58 analogs AZ5994 and AZ7004, compounds that in addition to having allosteric FFAR2 modulating effects, also resemble the effect of an orthosteric agonist. The precise receptor sites involved in the modulation are not known, but the fact that one modulator turns the other into a potent neutrophil activating agonist, suggests that the two interact with different allosteric binding sites. The signals generated by the orthosteric agonist propionate in neutrophils sensitized by the allosteric FFAR2s modulated by AZ1729 or Cmp58, are balanced in the sense that the activation of the superoxide producing NADPH-oxidase is accompanied by a transient rise in [Ca^2+^]_i_. Although the allosteric GPCR modulation concept is fairly new, it has rapidly developed into a very active field of research with the ultimate goal to develop clinically useful therapeutics (23). The fact that allosteric modulators may bind to several sites that are distinct from the orthosteric binding sites, but largely affect the functions induced by orthosteric agonists, raises questions about natural occurring endogenous regulators of GPCR functions (24). Our knowledge about the mechanisms for allosteric GPCR modulation, will be of help in the search for, and characterization of, natural allosteric modulators with regulatory functions, and this field of research will eventually turn from being a laboratory curiosity to an area of research that have direct clinical relevance. An increased understanding of how neutrophil GPCR activities are regulated may, thus, be of direct importance both in physiological and pathological settings, and might enable the development of novel prophylactic, treatment strategies and/or secondary prevention of inflammatory diseases.

To conclude, we describe novel activation mechanisms for the G-protein coupled free fatty acid receptor 2 (FFAR2). The data presented also provide direct evidence for biased FFAR2 signaling, resulting in an activation of the neutrophil NADPH-oxidase without involvement of the PLC-PIP_2_-IP_3_-Ca^2+^-route. The terminology for compounds that affect signaling and/or functional response induced by on orthosteric agonist is complex, but even if it has been suggested that ago-allosteric agonist/modulators in contrast to ordinary allosteric enhancers improve the maximal efficacy of the orthosteric agonist, the basic function of the two types of enhancers is similar (25). It is clear that both AZ5994 and AZ7004 have the dual functions acting as both a direct activating agonist and as an allosteric enhancer, suggesting that they should be classified as an allosteric modulating agonist. More importantly, i) the allosteric effect is achieved through binding to the same binding site as Cmp58 as illustrated by the fact that neutrophils are activated when AZ5994 and AZ7004 are combined with AZ1729 as well as with “AZ1729-like” allosteric modulators but not when combined with Cmp58, ii) the activation profile of AZ5994/AZ7004 differs from that of Cmp58 in that the signals generated when combined with AZ1729 initiate also a transient rise in [Ca^2+^]_i_, and iii) the propionate response is increased also when AZ5994/AZ7004 have been used as the activating/sensitizing agonist. The novel neutrophil activation patterns and receptor down-stream signaling mediated by two cross-sensitizing allosteric modulators represent a new regulatory mechanism that controls FFAR2 function.

## Experimental procedures

### Chemicals

Isoluminol, TNF-a, ATP, propionic acid, and bovine serum albumin (BSA), were purchased from Sigma (Sigma Chemical Co., St. Louis, MO, USA). Dextran and Ficoll-Paque were obtained from GE-Healthcare Bio-Science (Uppsala, Sweden). Fura 2-AM was from Molecular Probes/Thermo Fisher Scientific (Waltham, MA, USA), and horseradish peroxidase (HRP) was obtained from Boehringer Mannheim (Mannheim, Germany). The allosteric FFAR2 modulator Cmp58 ((*S*)-2-(4-chlorophenyl)-3,3-dimethyl-*N*-(5-phenylthiazol-2-yl)butanamide was obtained from Calbiochem-Merck Millipore (Billerica, USA) and TOCRIS (Bristol, UK). The FFAR2 agonist compound 1 (Cmp1; (*R*)-3-benzyl-4-(cyclopropyl(4-(2,5-dichlorophenyl)-thiazol-2-yl)amino)-4-oxobutanoic acid) and the antagonist CATPB ((*S*)-3-(2-(3-chlorophenyl)acetamido)-4-(4-(trifluoromethyl)phenyl) butanoic acid synthesized as described previously (15,26,27), were obtained (as generous gifts) from Trond Ulven (Odense University, Denmark). The allosteric FFAR2 modulator AZ1729 (10,12) together with all the other compounds included in the study (structures are shown in Fig 3A and B) were provided by AstraZeneca (Gothenburg, Sweden).

Subsequent dilutions of receptor ligand and other reagents were made in Krebs-Ringer Glucose phosphate buffer (KRG; 120 mM NaCl, 4.9 mM KCl, 1.7 mM KH_2_PO_4_, 8.3 mM NaH_2_PO_4_, 1.2 mM MgSO_4_, 10 mM glucose, and 1 mM CaCl_2_ in dH_2_O, pH 7.3).

### Isolation of human neutrophils

Neutrophils were isolated from buffy coats from healthy blood donors by dextran sedimentation and Ficoll-Paque gradient centrifugation as described by Bøyum (28). Remaining erythrocytes were removed by hypotonic lysis and the neutrophils were washed and resuspended in KRG. More than 90 percent of the cells were neutrophils and routinely with a viability of >95%. To amplify the activation signals the neutrophils were primed with TNF-a (10 ng/mL for 20 min at 37°C), and then stored on ice until use.

### Measuring NADPH-oxidase activity

Isoluminol-enhanced chemiluminescence (CL) technique was used to measure superoxide production, the precursor of production of reactive oxygen species (ROS), by the NADPH-oxidase activity as described (29,30). In short, the measurements were performed in a six-channel Biolumat LB 9505 (Berthold Co., Wildbad, Germany), using disposable 4-mL polypropylene tubes and a 900 μL reaction mixture containing 10^5^ neutrophils, isoluminol (0.2 μM) and HRP (4 Units/mL). The tubes were equilibrated for 5 min at 37°C, before addition of agonist (100 μL) and the light emission was recorded continuously over time. In experiments where the effects of receptor specific antagonists were determined, these were added to the reaction mixture 1–5 min before stimulation with control neutrophils incubated under the same condition but in the absence of antagonist run in parallel for comparison.

### Calcium mobilization

Neutrophils at a density of 1–3×10^6^ cells/mL were washed with Ca^2+^-free KRG and centrifuged at 220 *g*. The cell pellets were re-suspended at a density of 2×10^7^ cells/mL in KRG containing 0.1% BSA, and loaded with 2 μM FURA 2-AM for 30 min at room temperature. The cells were then washed once with KRG and resuspended in the same buffer at a density of 2×10^7^ cells/mL. Calcium measurements were carried out in a Perkin Elmer fluorescence spectrophotometer (LC50), with excitation wavelengths of 340 nm and 380 nm, an emission wavelength of 509 nm, and slit widths of 5 nm and 10 nm, respectively. The transient rise in intracellular calcium is presented as the ratio of fluorescence intensities (340 nm / 380 nm) detected.

### Analysis

A two-dimensional structural similarity map between the compounds included in the study and the suggested FFAR2 classification was based on extended-connectivity fingerprints (ECFP_6) were used to calculate Tanimoto scores and generate a two-dimensional structural similarity map (Rogers D. Extended-Connectivity Fingerprints *(31)*.

## Ethics Statement

In this study, conducted at the Sahlgrenska Academy in Sweden, buffy coats obtained from the blood bank at Sahlgrenska University Hospital, Gothenburg, Sweden have been used. According to the Swedish legislation section code 4§ 3p SFS 2003:460 (Lag om etikprövning av forskning som avser människor), no ethical approval was needed since the buffy coats were provided anonymously and could not be traced back to a specific donor.

## Data availability

All data are contained within the article.

## Acknowledgements

We thank the members of the Phagocyte Research Group at the Sahlgrenska Academy, University of Gothenburg, for critically discussing the results and the manuscript.

## Authorship contributions

*Participation in research design:* Lind, Holdfeldt, Mårtensson,

*Conducted experiments:* Lind, Holdfeldt,

*Contributed chemical tools and performed data analysis:* Granberg

*Performed data analysis:* Lind, Holdfeldt, Granberg, Forsman, Dahlgren

*Planned and supervised the research and wrote the manuscript:* Forsman, Dahlgren

*Contributed to the writing of the manuscript:* Lind, Holdfeldt, Mårtensson, Granberg

